# Rapid spread of knockdown resistance allele frequencies in a suburban population of *Aedes albopictus*

**DOI:** 10.1101/2025.06.30.662396

**Authors:** Jennifer F. Baltzegar, Cole D. Butler, Jessica Y. Ding, Chay M. Beeson, Emily MX Reed, Michael H. Reiskind, Martha O. Burford Reiskind

## Abstract

*Aedes albopictus* is a major vector of arboviral diseases and is often targeted by pyrethroid-based mosquito control in residential areas. While knockdown resistance (*kdr*) mutations are well documented in *Ae. aegypti*, their emergence in *Ae. albopictus* has been less studied, particularly in suburban environments where insecticide application is often uncoordinated. Understanding the temporal and spatial dynamics of resistance evolution in this context is critical for preserving the effectiveness of public health interventions. We conducted longitudinal sampling of *Ae. albopictus* populations in Wake County, North Carolina, from 2016 to 2024. Using a novel allele-specific PCR melt curve assay, we genotyped 2,273 mosquitoes at the F1534S locus in the voltage-gated sodium channel gene. Resistance allele frequencies were calculated annually and mapped across the county for three key years—2016, 2018, and 2023— representing the pre-emergence, initial detection, and widespread phases of resistance development. Selection and dominance were estimated using the Wright-Fisher Approximate Bayesian Computation algorithm for locus F1534S. The F1534S resistance allele was first detected in 2018 at a central, affluent neighborhood. By 2023, the allele had spread throughout the sampling region, with the highest frequencies near the site of first detection. Resistance allele frequency peaked at 0.36 in 2023, accompanied by an increase in heterozygous and homozygous resistance genotypes. Temporally sampled sites showed consistent trends in rising resistance, with all temporally-sampled locations harboring resistance genotypes by 2022. The resistance allele was found to be under high selection pressure and partially recessive in this population.

Our findings reveal rapid emergence and spatial expansion of the F1534S *kdr* allele in *Ae. albopictus* populations in a suburban setting. The pattern of spread suggests strong local selection pressure, likely driven by residential pyrethroid use. These results highlight the need for proactive resistance monitoring and integrated management strategies that address private-sector contributions to insecticide selection pressure.

## Introduction

Insecticide resistance is a global public health challenge and a major threat to the control of vector-borne diseases (1). In mosquitoes, pyrethroids are the most widely used class of insecticides as their relatively low cost and low mammalian toxicity makes them one of the safest options for use in and around homes (2,3). With continued use, resistance to pyrethroids can evolve rapidly, undermining the efficacy of these compounds and compromising both epidemic response and nuisance mosquito control. It is critical that best practices for insecticide resistance management are informed by an understanding of how resistance emerges and spreads, particularly in systems, such as those found across much of the United States, where centralized vector control programs are limited or absent.

*Aedes albopictus*, a mosquito species native to Asia, now inhabits every continent except Antarctica and Australia. Its expanding range is fueled by globalization, trade, and a warming climate that opens new habitats for this temperate-adapted species (4). *Ae. albopictus* is a competent vector for dengue, chikungunya, Zika, and other arboviruses, and is a major vector of dog heartworm (4). In addition to its role in disease transmission, it is a major nuisance mosquito, particularly in suburban areas, where its aggressive daytime biting drives high demand for control (5).

In the United States, efforts to control *Ae. albopictus* are often decentralized. In many suburban settings, including Wake County, North Carolina, mosquito control decisions are typically determined by homeowners rather than through publicly funded municipal programs. In the absence of coordinated public health infrastructure, homeowners contract private companies that offer seasonal pyrethroid-based adulticide treatments often with repeated applications. This uncoordinated approach creates localized and repeated selection pressure for resistance to build yet rarely includes resistance monitoring or systematic management practices.

Sustained pyrethroid exposure can lead to full phenotypic resistance in mosquito populations. Mutations in the voltage-gated sodium channel (*vgsc*) gene, the primary target of pyrethroids, are a well-documented mechanism of knockdown resistance (*kdr*) and can evolve rapidly in response to insecticide pressure (6). While most studies of *kdr* mutations in *Ae. albopictus* have focused on urban environments with centralized control (7–10), suburban landscapes present a unique opportunity to study resistance evolution in a fragmented, free-market system. In these settings, variation in application frequency, treatment timing, and coverage likely leads to heterogeneous selection pressures across the landscape.

The F1534S mutation in the *vgsc* gene is the most commonly reported *kdr* allele in *Ae. albopictus*. It was first identified in populations from China, where it has been linked to phenotypic resistance to pyrethroids (10). It has now been detected in *Ae. albopictus* populations on multiple continents, including in the United States (9). However, its emergence and spread have not been studied in the context of a suburban environment characterized by variable insecticide exposure.

In this study, we develop a high-throughput, cost-effective allele-specific PCR (AS-PCR) assay to rapidly genotype individual *Ae. albopictus* mosquitoes at the F1534S locus. We apply this assay to mosquito collections spanning nine years (2016–2024) across Wake County, North Carolina, to track temporal and spatial patterns of resistance. With our temporally repeated sampling, we also evaluate the selection pressure in this environment and assess the dominance of the resistance allele. Our goal is to document how quickly resistance develops in a decentralized control landscape and provide context for vector control and resistance management.

## Methods

### Sample collections

#### Temporal Sampling (Five sites)

We collected *Ae. albopictus* adults at five sites starting in 2016 and continuing to 2024, as part of on-going mosquito surveillance in Wake County, NC (Fig 1). These five sites are located in suburban areas, two are commercial, one is institutional, located at North Carolina State University, and two are at residential homes. We collected adults weekly every summer using BG-Sentinel traps, and in 2016, we used ovitraps to collect eggs (11,12).

**Fig 1.**
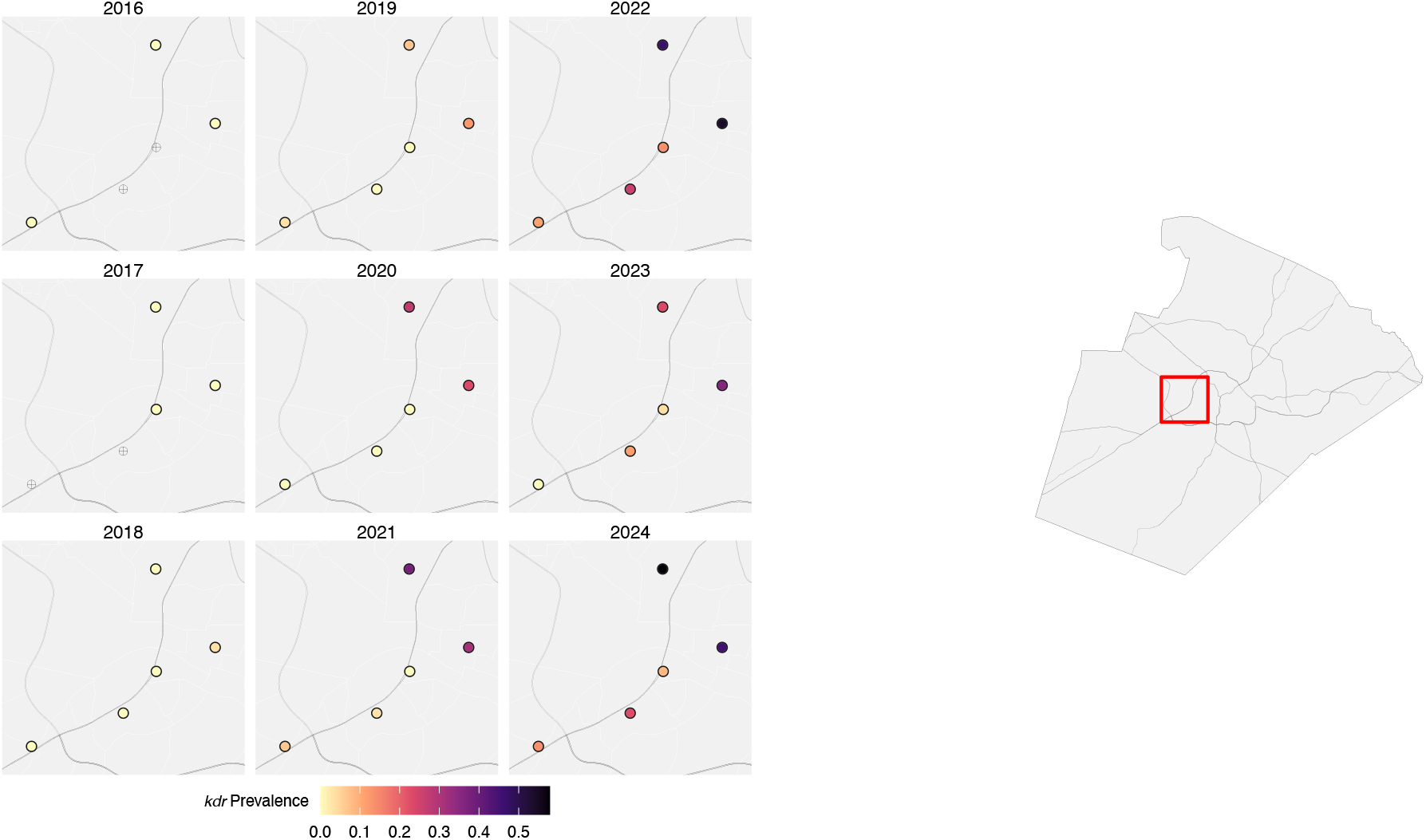
Prevalence of kdr genotypes at frequently sampled locations within Wake County. Each circle represents a single sample location (i.e., homeowner’s property) within Wake County, North Carolina. The color of the circle represents the allele frequency of kdr at that sample location. Lighter colors indicate lower kdr frequencies while darker colors represent higher kdr frequencies. No mosquitoes contained kdr alleles in 2016 and 2017. Resistance first emerged within these samples in 2018 and quickly rose to very high frequencies in some sites. As of 2024, all five of the frequently sampled locations have moderate to high levels of F1534S frequencies.

#### Wide Spatial Sampling (2016, 2018, 2023)

We conducted spatial sampling for population genetic and ecological studies in 2016 and 2018 that spanned an urban rural gradient. We collected *Ae. albopictus* eggs from sixteen sites in 2016 weekly between 15 April – 26 October 2016 as part of a state-wide mosquito survey effort to characterize the distribution and abundance of *Aedes* species in North Carolina (11,13). Using ovitraps, we collected eggs which were later hatched, reared, and identified (for full methods, see (13)). In 2018, we collected *Ae. albopictus* adults from 52 sites using BG-Sentinel traps as part of a population genetic study (11) (Fig 2). We chose the five sites that were part of the annual monitoring program, and the remaining sites were chosen randomly (for full methods, see (11)). These sites spanned an urban to rural gradient in Wake County, NC. Mosquitoes were collected weekly from 15 April – 26 October 2016 and again from 7 June – 25 July 2018, leaving the traps out to collect mosquitoes for 24 hours (11,12). The larger spatial sampling in 2016 and 2018 allowed us to highlight where and when F1534S first appeared and how the mutation spread. In 2023, we conducted an *a priori-*designed landscape study at 40 houses across 31 blocks across a wealth and neighborhood age gradient, in addition to the regularly sampled five sites (Fig 2)(14). Adult *Ae. albopictus* were collected using BG-Sentinel traps. Samples from all spatial and temporal studies were stored by date and location in a -80°C freezer for future processing.

**Fig 2.**
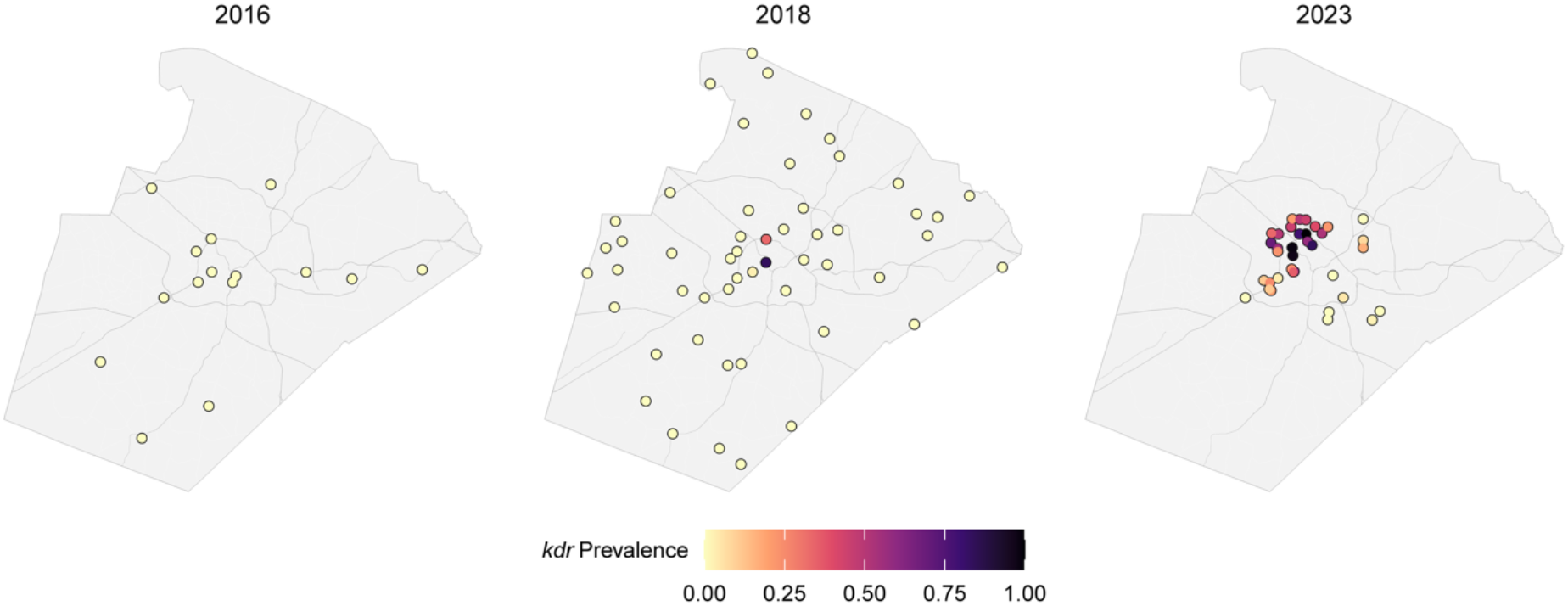
Prevalence of kdr genotypes in Wake County. Each circle represents a single mosquito collection site within Wake County, North Carolina. The color of the circle represents the allele frequency of kdr resistance at that sample location. Lighter colors indicate lower kdr frequencies and darker colors represent higher kdr frequencies.

### DNA Isolation and Quantification

Genomic DNA was extracted from individual mosquitoes using the Qiagen DNeasy Blood and Tissue Kit, with modifications to the manufacturer’s protocol for insect specimens. The head and thorax of each mosquito were homogenized via bead disruption in ATL buffer and proteinase K. Samples were incubated overnight at 56°C. On the following day, wash steps were performed as prescribed by the protocol, followed by two elution steps in warmed nuclease-free water, yielding a final elution volume of 100 µL. DNA concentration was quantified using a Qubit Fluorometer (ThermoFisher Scientific) with the Qubit 1X dsDNA HS Assay Kit (Cat. No. Q33231).

### Allele-specific PCR genotyping

To genotype individual mosquitoes at the *kdr* locus F1534S, we developed an allele-specific PCR (AS-PCR) melt curve assay based on methods previously used for *Ae. aegypti* (6,15,16).

Three primers were designed: one common reverse primer (F1534-Rev: TCTGCTCGTTGAAGTTGTCGAT) and two allele-specific forward primers. The forward primers were designed to uniquely amplify either the susceptible allele (F1534-Fwd: GCGGGCTCTACTTCGTGTTCTTCATCATATT) or the resistance allele (S1534-Fwd: GCGGGCAGGGCGGCGGGGGCGGGGCCTCTACTTCGTGTTCTTCATCATGTC), with GC-rich tags of differing lengths to ensure distinct melting temperatures (∼4°C apart) of the resulting PCR products.

Each 10 µL PCR reaction contained: 5.0 µL dH_2_O, 0.2 µL S1534-Fwd primer, 0.4 µL F1534-Fwd primer, 0.4 µL F1534-Rev primer, 3.0 µL PerfeCTa SYBR Green SuperMix for iQ (VWR: 101414-144), and 1.0 µL gDNA. Reactions were performed on a Bio-Rad C1000 Touch Thermal Cycler CFX384 Real-Time System (Genomic Sciences Laboratory, North Carolina State University) under the following cycling conditions: initial denaturation at 95°C for 2 min; 35 cycles of 95°C for 30 sec, 58°C for 30 sec, and 72°C for 30 sec; followed by a final extension at 72°C for 2 min. A melt curve analysis was then conducted from 65°C to 95°C in 0.2°C increments every 10 seconds, with fluorescence measured at each step.

Melting temperature (Tm) peaks were visualized using Bio-Rad Maestro software. The susceptible and resistance alleles produced Tm peaks at approximately 78°C and 82°C, respectively. Accordingly, homozygous susceptible individuals showed a single peak at 78°C, homozygous resistant individuals showed a single peak at 82°C, and heterozygotes displayed both peaks. To increase confidence in our allele estimates, each sample was genotyped in duplicate using the AS-PCR assay, and only those with matching replicate results were included in downstream analyses.

### Measures of genetic estimates

Consensus genotypes were determined by comparing results from the two replicate AS-PCR assays for each sample. Samples that failed to amplify in both replicates or produced discordant results were excluded from further analysis. For the remaining samples, genotype and allele frequencies were calculated for each collection year using a custom R script (17). The selection coefficient and dominance of F1534S were estimated by employing the Wright-Fisher approximate Bayesian calculation (WF-ABC) model (18). Hardy-Weinberg equilibrium (HWE) and linkage estimates were calculated in the R using the *genetics* package (19). All maps were generated using custom R scripts and included annual allele frequency per site.

## Results

### Temporal trends in kdr allele frequency

Between 2016 and 2024, we genotyped 2,669 *Ae. albopictus* mosquitoes from 114 unique collection sites in Wake County, North Carolina using allele-specific PCR targeting the F1534S *kdr* mutation. Only samples with corroborating genotypes across two technical replicates were retained for analysis. Resistance alleles were not detected in the first two years of the study (2016–2017), but their frequency increased steadily thereafter, reaching a maximum of 0.36 (95% CI ± 0.027) in 2023 (Table 1). In 2024, the resistance allele frequency declined to 0.25 (95% CI ± 0.044), perhaps due to fewer sites sampled in a heterogenous landscape. Across the same period, the proportion of homozygous susceptible genotypes (SS) decreased, while heterozygous (SR) and homozygous resistant (RR) genotypes increased.

**Table 1.** Genotype and allele frequency results by year. Frequency of each genotype recorded by year for F1534S locus in Aedes albopictus collected from Wake County, North Carolina from 2016 to 2024. The genotypes are SS (homozygous susceptible), SR (heterozygotes, RR (homozygous resistant).

| Year | Sites (n) | Mosquitoes (n) | Freq R Allele | Genotype Frequencies |  |  |
| --- | --- | --- | --- | --- | --- | --- |
|  |  |  |  | SS | SR | RR |
| 2016 | 16 | 225 | 0.00 | 1.00 | 0.00 | 0.00 |
| 2017 | 3 | 82 | 0.00 | 1.00 | 0.00 | 0.00 |
| 2018 | 52 | 516 | 0.01 | 0.98 | 0.00 | 0.01 |
| 2019 | 5 | 154 | 0.05 | 0.93 | 0.05 | 0.02 |
| 2020 | 5 | 153 | 0.10 | 0.86 | 0.08 | 0.06 |
| 2021 | 5 | 173 | 0.16 | 0.77 | 0.13 | 0.10 |
| 2022 | 5 | 143 | 0.29 | 0.61 | 0.20 | 0.20 |
| 2023 | 36 | 609 | 0.36 | 0.52 | 0.25 | 0.23 |
| 2024 | 5 | 218 | 0.25 | 0.67 | 0.14 | 0.18 |

### Temporal dynamics of resistance

To track resistance emergence and spread at consistent locations, we conducted repeated sampling at five fixed sites, within suburban residential areas, from 2016 to 2024. In 2016 and 2017, we only found mosquitoes at three of the five sites and the mosquitoes collected showed only homozygous susceptible genotypes. The F1534S resistance allele first appeared in 2018 at one of the sampled sites. (A nearby second site included in the expanded spatial sampling for 2018 (see below) also tested positive for resistance alleles in 2018 (Figs. 1 and 2)). By 2022, all five of the temporally sampled sites tested positive for mosquitoes carrying at least one copy of the resistance allele (Fig 1). By 2024, each site exhibited moderate to high frequencies of F1534S, indicating rapid and sustained increases in resistance allele frequency, potentially driven by local selection, migration, or both.

### Spatial expansion of resistance

Broader spatial sampling conducted in 2016, 2018, and 2023 captured a wider geographic pattern of *kdr* allele distribution across Wake County. Following a similar pattern to resistance emergence as the temporally sampled sites, all genotyped mosquitoes were homozygous susceptible in 2016. In 2018, the resistance allele was detected at low frequency and localized primarily at two sites near the center of the county (including one of the temporally sampled sites), an area characterized by older and more affluent neighborhoods. By 2023, resistance alleles were found throughout the sampling region, with the highest allele frequencies concentrated near the original 2018 detection zone (Fig 2). Peripheral areas of the county generally exhibited lower resistance frequencies.

### Genetic parameters of resistance

Using time series allele frequency data collected from both the temporal and spatial datasets, we estimated the posterior distribution of the selection coefficient (s) and dominance (h) of the F1534S resistance allele using the WFABC algorithm (18). The mean selection coefficient was 0.68, with a 95% credible interval of 0.25 to 0.99 (2.5^th^ to 97.5^th^ quantile). The mean dominance coefficient was 0.27, with a 95% credible interval of 0.01 to 0.71, indicating that the resistance allele is partially recessive. Box plots summarizing the marginal posterior distributions of s and h (Fig 3) highlight estimate uncertainty and inter-correlation. However, the two-dimensional kernel density plot of the joint posterior distributions (Fig 4) supports a region of highest posterior density characterized by strong selection and partial recessivity.

**Fig 3.**
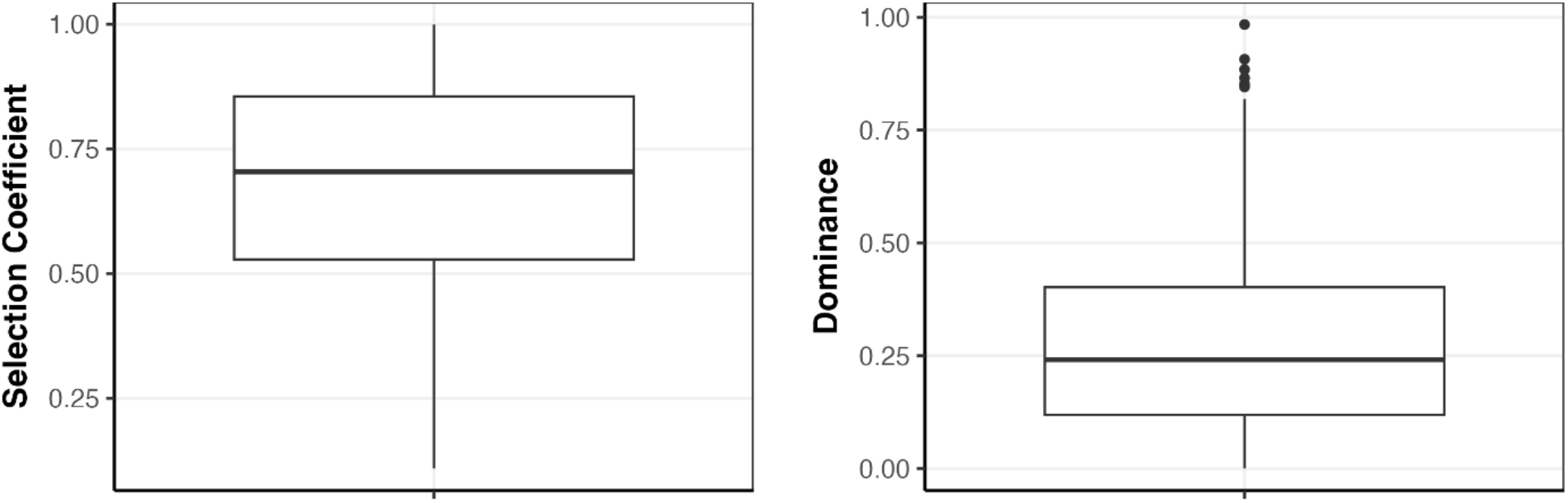
Marginal posterior distribution estimates of the selection coeVicient (s) and dominance (h) for the F1534S allele. Box plots show the central tendency and spread of posterior samples, indicating strong selection (mean = 0.68) and partial recessivity (mean = 0.27).

**Fig 4.**
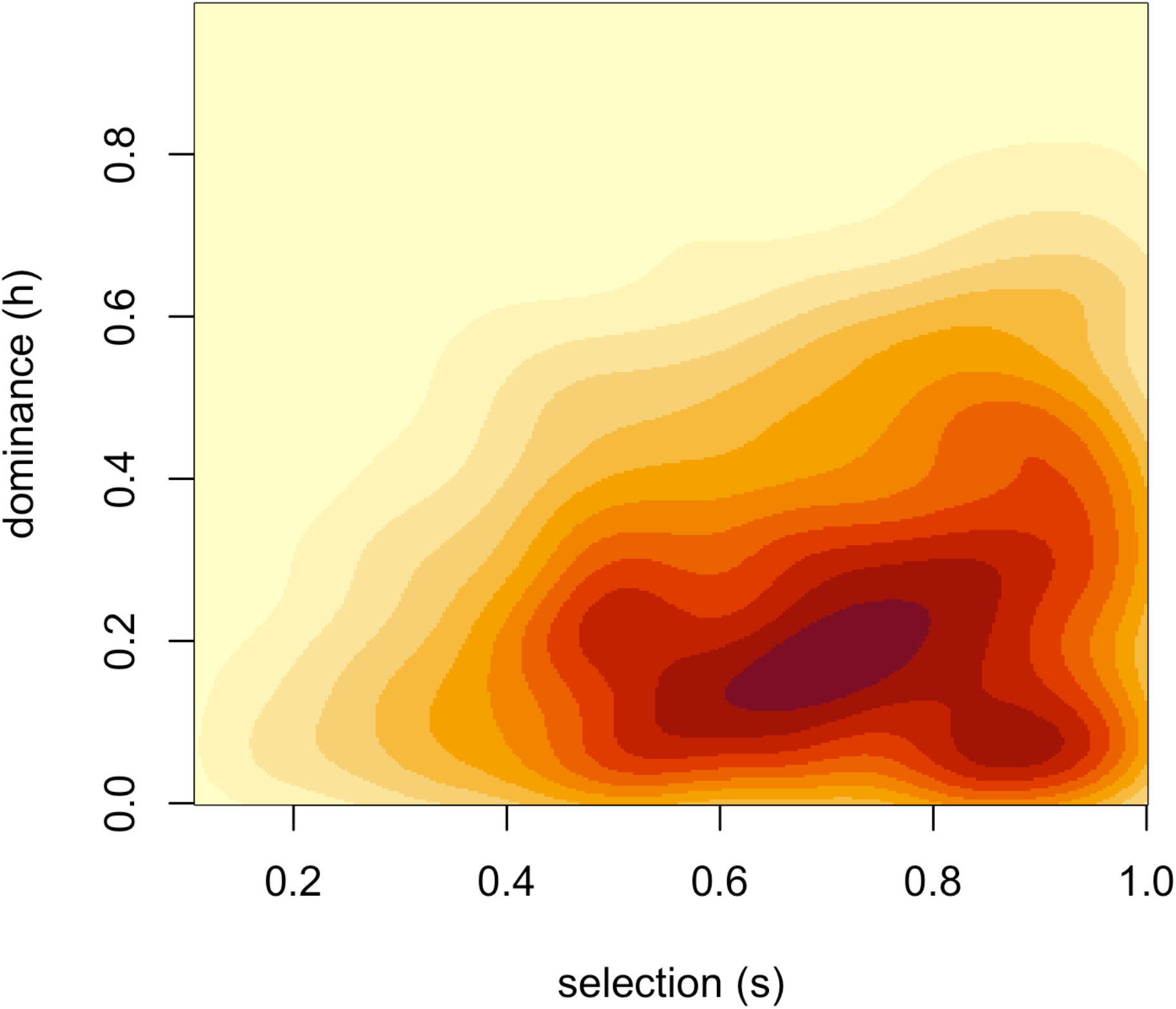
Two-dimensional kernel density plot of the joint posterior distributions of the selection coeVicient (s) and dominance (h), with darker colors representing regions of highest posterior density.

Genotype frequencies at the F1534S locus deviated significantly from Hardy-Weinberg expectations (exact test, *p* < 2.2 × 10^−16^). The observed heterozygote frequency (R/S = 5.6%) was substantially lower than that expected under HWE (13.7%), resulting in a significant excess of homozygotes. The observed overall disequilibrium (D = -0.081) and strong negatively scaled disequilibrium (D’ = -3.02, 95% CI: -3.37 to -2.72) also indicate a significant deficit of heterozygotes, consistent with ongoing selection at this locus. This significant heterozygote deficit may reflect a fitness advantage of homozygous resistant individuals undergoing selection, consistent with a partially or fully recessive resistance allele.

## Discussion

Monitoring insecticide resistance in mosquito populations is critical for maintaining effective vector control strategies. In this study, we provide the first longitudinal dataset documenting the emergence and spread of the F1534S *kdr* allele in *Aedes albopictus* from a suburban environment in the United States. Over the nine-year study period, we observed a clear increase in both the frequency and geographic distribution of this resistance allele in Wake County, North Carolina, consistent with strong and ongoing selection pressure.

Our results indicate that the resistance allele at locus F1534S (*i*.*e*., Ser1534) was absent from local *Ae. albopictus* populations until 2018. Since its emergence, it has risen in frequency across all temporally sampled sites, reaching moderate to high frequency by 2023. Spatial analysis further revealed that the resistance allele first appeared near the geographic center of Wake County, an area characterized by older, more affluent neighborhoods. By 2023, Ser1534 was detected at high frequencies across much of the county, suggesting widespread establishment. The resistance allele frequency was slightly lower in 2024 compared to 2023. This decline is likely an artifact of reduced sampling effort and fewer collection sites in 2024 (n = 5) compared to 2023 (n = 36), rather than a true reversal in resistance trends. Future work should maintain consistent spatial coverage across years to better track ongoing dynamics. These results demonstrate not only the rapid spread of resistance but also a potentially directional pattern of dissemination, likely reflecting human-mediated selection and local mosquito dispersal dynamics.

We hypothesize that residential insecticide use, particularly through professionally applied barrier sprays by private pest control companies, is a major driver of this resistance evolution. The observed pattern of initial emergence in wealthier neighborhoods aligns with previous studies linking socioeconomic status and insecticide use frequency to elevated selection pressure on mosquito populations in *Aedes aegypti* and our forthcoming work on *Ae. albopictus* (14,20). Although *Ae. albopictus* in Wake County has not yet been implicated in major arboviral outbreaks, the increasing risk of diseases such as dengue and chikungunya in the southeastern United States highlights the importance of preserving effective vector control tools (21).

Non-synonymous mutations at amino acid position 1534 have previously been identified as a key marker of pyrethroid resistance in *Ae. albopictus* (F1534S) and *Ae. aegypti* (F1534C) (7–10). In both species, point mutations at this locus often represent the first step in the evolution of pyrethroid resistance (6,7). While additional mechanisms such as metabolic detoxification contribute to the complete pyrethroid resistance phenotype, Ser1534 is associated with reduced sensitivity to pyrethroids and can serve as a sentinel indicator of resistance evolution (10,22).

Our results, together with previous work in this population, support the role of Ser1534 as a sentinel allele and early indicator of increasing insecticide resistance (23). We observed a significant departure from Hardy-Weinberg equilibrium at this locus, consistent with ongoing selection for the resistance allele. The estimated selection coefficient for Ser1534 was very high (s = 0.68), indicating strong directional selection in favor of the resistance allele. This is likely driven by continued insecticide pressure from homeowner use of commercial pyrethroid products, even in the absence of organized mosquito control.

Given its partial recessivity (h = 0.27), the allele frequency of Ser1534 is expected to increase rapidly once homozygotes become common, given that heterozygotes retain partial susceptibility. This would allow the wild-type allele to persist under moderate selection pressure, thus providing an opportunity to preserve insecticide efficacy (24). These dynamics have important implications for resistance management as incomplete dominance can delay the fixation of the resistance allele, allowing for low-level resistance to persist undetected – in the absence of molecular genotyping – until phenotypic resistance becomes widespread and control failure occurs.

Our findings underscore the urgent need for routine resistance monitoring in *Ae. albopictus*, particularly in suburban settings where private pesticide use is common. We recommend that vector control programs incorporate genetic surveillance of locus F1534S and other *kdr* mutations into their operations. Furthermore, public health authorities and pest control companies should work collaboratively to improve and promote integrated vector management strategies that slow the spread of phenotypic resistance to insecticides.

## Conclusion

Our study documents the rapid emergence and spread of the F1534S *kdr* resistance allele in *Aedes albopictus* across Wake County, North Carolina over a nine-year period. The resistance allele was first detected in 2018 and has since increased in frequency and geographic extent, likely driven by selection pressure from suburban residential use of pyrethroid-based insecticides. Given the persistence and potential public health impact of this mutation, we recommend routine monitoring of *kdr* alleles in *Ae. albopictus* and greater collaboration between public health agencies and private pest control services to implement improved insecticide resistance management strategies.

## Acknowledgements

We would like to thank the Genetics and Genomic Academy and the Global One Health Academy at North Carolina State University for supporting this work.

## Notes

### Competing Interest Statement

The authors have declared no competing interest.

## References

1. Dusfour I, Vontas J, David JP, Weetman D, Fonseca DM, Corbel V, et al. Management of insecticide resistance in the major Aedes vectors of arboviruses: Advances and challenges. PLoS Negl Trop Dis. 2019 Oct 10;13(10):e0007615.

2. Liu N, Xu Q, Zhu F, Zhang L. Pyrethroid resistance in mosquitoes. Insect Sci. 2006 Jun;13(3):159–66.

3. Gajendiran A, Abraham J. An overview of pyrethroid insecticides. Front Biol. 2018 Apr;13(2):79–90.

4. Bonizzoni M, Gasperi G, Chen X, James AA. The invasive mosquito species Aedes albopictus: current knowledge and future perspectives. Trends Parasitol. 2013 Sep;29(9):460–8.

5. Paupy C, Delatte H, Bagny L, Corbel V, Fontenille D. Aedes albopictus, an arbovirus vector: From the darkness to the light. Microbes Infect. 2009 Dec;11(14–15):1177–85.

6. Baltzegar J, Vella M, Gunning C, Vasquez G, Astete H, Stell F, et al. Rapid evolution of knockdown resistance haplotypes in response to pyrethroid selection in Aedes aegypti. Evol Appl. 2021 Aug;14(8):2098–113.

7. Zhang Y, Wang D, Shi W, Zhou J, Xiang Y, Guan Y, et al. Resistance to pyrethroids and the relationship between adult resistance and knockdown resistance (kdr) mutations in Aedes albopictus in dengue surveillance areas of Guizhou Province, China. Sci Rep. 2024 May 28;14(1):12216.

8. Wu Y, Liu Q, Qi Y, Wu Y, Ni Q, Chen W, et al. Knockdown Resistance (kdr) Mutations I1532T and F1534S Were Identified in Aedes albopictus Field Populations in Zhejiang Province, Central China. Front Cell Infect Microbiol. 2021 Jun 29;11:702081.

9. Xu J, Bonizzoni M, Zhong D, Zhou G, Cai S, Li Y, et al. Multi-country Survey Revealed Prevalent and Novel F1534S Mutation in Voltage-Gated Sodium Channel (VGSC) Gene in Aedes albopictus. Kittayapong P, editor. PLoS Negl Trop Dis. 2016 May 4;10(5):e0004696.

10. Chen H, Li K, Wang X, Yang X, Lin Y, Cai F, et al. First identification of kdr allele F1534S in VGSC gene and its association with resistance to pyrethroid insecticides in Aedes albopictus populations from Haikou City, Hainan Island, China. Infect Dis Poverty. 2016 Dec;5(1):31.

11. Reed EM, Reiskind MH, Burford Reiskind MO. Spatiotemporal variation in abundance and genetic structure across the urban-rural landscape gradient: Aedes albopictus (Skuse, 1894) (Diptera: Culicidae) in Wake County, NC [Internet]. Genetics; 2025 [cited 2025 Jun 18]. Available from: http://biorxiv.org/lookup/doi/10.1101/2025.06.12.659333

12. Reed EMX, Reiskind MH, Burford Reiskind MO. Life-history stage and the population genetics of the tiger mosquito Aedes albopictus at a fine spatial scale. Med Vet Entomol. 2023 Mar;37(1):132–42.

13. Reed EMX, Byrd BD, Richards SL, Eckardt M, Williams C, Reiskind MH. A Statewide Survey of Container Aedes Mosquitoes (Diptera: Culicidae) in North Carolina, 2016:A Multiagency Surveillance Response to Zika Using Ovitraps. J Med Entomol. 2019 Feb 25;56(2):483–90.

14. Butler C, Ding J, Baltzegar J, Brown ZS, Burford Reiskind M, Reiskind MH. Socioeconomic predictors of knockdown resistance in Aedes albopictus (Diptera: Culicidae). Prep. 2025;

15. Yanola J, Somboon P, Walton C, Nachaiwieng W, Somwang P, Prapanthadara Laied. High-throughput assays for detection of the F1534C mutation in the voltage-gated sodium channel gene in permethrin-resistant Aedes aegypti and the distribution of this mutation throughout Thailand. Trop Med Int Health TM IH. 2011 Apr;16(4):501–9.

16. Deming R, Manrique-Saide P, Medina Barreiro A, Cardeña EUK, Che-Mendoza A, Jones B, et al. Spatial variation of insecticide resistance in the dengue vector Aedes aegypti presents unique vector control challenges. Parasit Vectors. 2016 Feb 4;9(1):67.

17. R Core Team. R: A Language and Environment for Statistical Computing. R Found Stat Comput Vienna Austria [Internet]. 2025; Available from: <https://www.R-project.org/>

18. Foll M, Shim H, Jensen JD. WFABC : a W right– F isher ABC -based approach for inferring effective population sizes and selection coefficients from time-sampled data. Mol Ecol Resour. 2015 Jan;15(1):87–98.

19. Gregory Warnes, with contributions from Gregor Gorjanc, Friedrich Leisch, and Michael Man. genetics: Population Genetics [Internet]. 2002 [cited 2025 Jun 18]. p. 1.3.8.1.3. Available from: https://CRAN.R-project.org/package=genetics

20. Fay JV, Espinola SL, Boaglio MV, Blariza MJ, Lopez K, Zelaya F, et al. Pyrethroid genetic resistance in the dengue vector (Aedes aegypti) in Posadas, Argentina. Front Public Health. 2023 Apr 27;11:1166007.

21. Ryan SJ, Carlson CJ, Mordecai EA, Johnson LR. Global expansion and redistribution of Aedes-borne virus transmission risk with climate change. PLoS Negl Trop Dis. 2019 Mar 28;13(3):e0007213.

22. Yang X, Zhou Y, Sun Y, Liu J, Jiang D. Multiple insecticide resistance and associated mechanisms to volatile pyrethroid in an Aedes albopictus population collected in southern China. Pestic Biochem Physiol. 2021 May;174:104823.

23. Abernathy HA, Hollingsworth BD, Giandomenico DA, Moser KA, Juliano JJ, Bowman NM, et al. Prevalence of Knock-Down Resistance F1534S Mutations in Aedes albopictus (Skuse) (Diptera: Culicidae) in North Carolina. Healy K, editor. J Med Entomol. 2022 Jul 13;59(4):1363–7.

24. Guo Y, Hu K, Zhou J, Xie Z, Zhao Y, Zhao S, et al. The dynamics of deltamethrin resistance evolution in Aedes albopictus has an impact on fitness and dengue virus type-2 vectorial capacity. BMC Biol. 2023 Sep 13;21(1):194.

